# The human-specific *RPGR* isoform *RPGR^s14/15^* and the clinically approved Rho/ROCK inhibitor Ripasudil represent therapeutic options to address *RPGR*-associated defects

**DOI:** 10.1101/2025.05.09.653106

**Authors:** Muhammad Usman, Paul Atigbire, Dennis Kastrati, Julia Milena Brinkhoff, Charlotte Luise Kluth, Jannis Marticke, Christoph Jüschke, John Neidhardt

**Affiliations:** Human Genetics, Medical Faculty-School of Medicine and Health Sciences, Carl von Ossietzky Universität Oldenburg, 26129 Oldenburg, Germany; Research Center Neurosensory Science, Carl von Ossietzky University Oldenburg, Oldenburg, Germany

**Author notes:** **Corresponding author:** John Neidhardt, Human Genetics, Faculty of Medicine and Health Sciences, University of Oldenburg, Carl-von-Ossietzky-Straße 9-11, 26129 Oldenburg, Germany.

**Keywords:** Retinitis Pigmentosa GTPase Regulator, CRISPR-Cas9, Isoform *RPGR^s14/15^*, Cilia Structure, Actin dynamics, F-actin, Ripasudil, Rho GTPases, ROCK kinases

## Abstract

Pathogenic variants in the *RPGR* gene are the primary cause of photoreceptor degeneration in X-linked retinitis pigmentosa (RP). Previous studies have linked *RPGR* dysfunction to defects in ciliary structure and actin turnover. *RPGR* encodes three major isoforms—*RPGR^1–19^*, *RPGR^ORF15^*, and the human-specific *RPGR^s14/15^*—yet the function of *RPGR^s14/15^* remains poorly understood. There is an urgent unmet need for effective treatments targeting *RPGR*-associated RP. We generated *RPGR* mutant hTERT-RPE1 cell lines and found that the loss of all *RPGR* isoforms resulted in pronounced ciliary defects, including aberrant cilia length and segmentation, along with disrupted actin turnover. Strikingly, cells expressing only the human-specific isoform *RPGR^s14/15^*closely resembled wild-type controls and were largely protected from these defects, underscoring a critical role for *RPGR^s14/15^* in maintaining ciliary integrity and actin dynamics. Furthermore, we demonstrated that pharmacological disruption of actin polymerization with cytochalasin D (CytoD) in control cells mimicked the ciliary abnormalities seen in *RPGR*_KO cells. Conversely, treatment with Ripasudil—a clinically approved Rho/ROCK inhibitor—rescued both ciliary and actin-related defects in *RPGR*_KO lines without observable side effects. In summary, our findings highlight the therapeutic relevance of the human-specific isoform *RPGR^s14/15^* and identify Ripasudil as a promising candidate for treating *RPGR*-associated retinal degeneration.

**Graphical Abstract:** 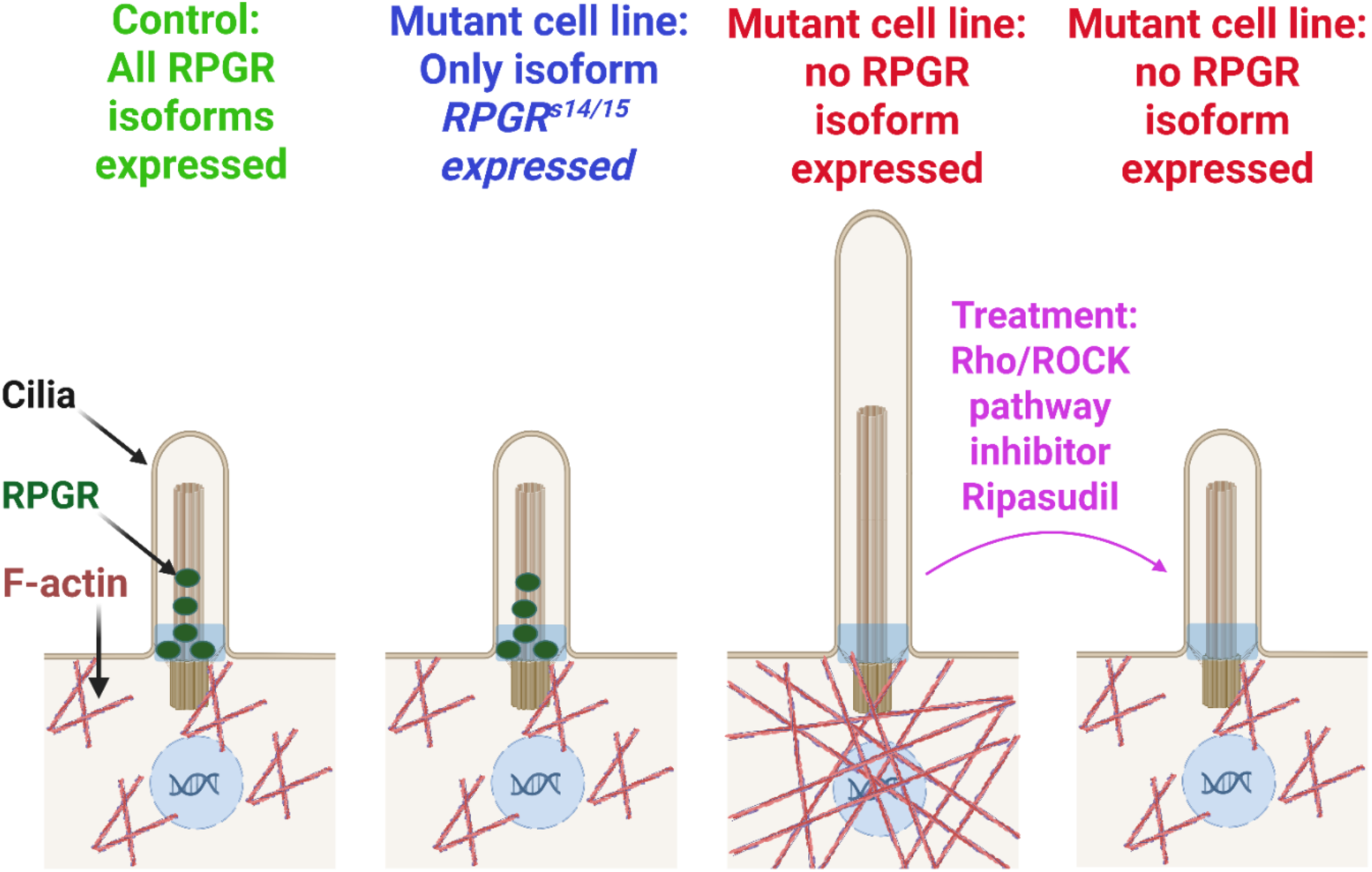

## Introduction

Primary cilia are evolutionarily conserved protrusions on the surface on many kinds of cells, including retinal cells ^1-5^. Historically, primary cilia were considered vestigial structures with minimal functional importance ^3^; however, more recent advancements demonstrated their importance in cellular signaling, tissue development, and homeostasis ^6-8^. Dysfunctional primary cilia have been associated with inherited diseases collectively termed ciliopathies, including retinitis pigmentosa (RP) ^9-11^. Structurally, primary cilia consist of the basal body (BB), that serves as the microtubule organizing center, the transition zone (TZ), which serves as a gatekeeper for ciliary trafficking, and the microtubule core axoneme, which acts as an antenna and hub for cellular signaling cascades ^12,13^. The axoneme of primary cilia has been compartmentalized into two major segments: The polyglutamylated proximal segment (PS) and the poorly glutamylated distal segment (DS). The ratio between the PS and DS length is crucial for ciliary functions ^14^.

Retinitis pigmentosa (RP, OMIM #268000) is a blinding disease caused by the progressive loss of rod and cone photoreceptors. It is estimated that 1 in every 4000 people globally is affected by RP. X-chromosomal RP (XLRP) is among the most severe forms of RP ^15^ and is frequently associated with genomic sequence alterations (mutations) in the Retinitis pigmentosa GTPase regulator (*RPGR [MIM312610]*) ^16-19^. The *RPGR* gene is located on the short arm of chromosome X, and mutations in the *RPGR* gene account for 12-15% of all RP patients, as well as 70-80% of the XLRP cases ^20-24^.

The RPGR protein localizes to the TZ of the primary cilium and to the connecting cilium of the photoreceptors ^25,26^. Loss of RPGR from the TZ of primary cilia causes ciliary length defects, which have been associated with the clinical manifestation of RP ^26,27^.

The human *RPGR* gene consists of 19 exons and encodes three major protein-coding isoforms, including the constitutive isoform *RPGR^1-19^*, isoform *RPGR^s14/15^*, and isoform *RPGR^ORF15^*. Isoform *RPGR^1-19^* includes all 19 exons, while isoform *RPGR^s14/15^* is generated by an in-frame skipping of exons 14 and 15. Isoform *RPGR^ORF15^* contains parts of intron 15 as an open reading frame and skips exons 16-19. Isoform *RPGR^1-19^*and *RPGR^s14/15^* encode proteins of 815 aa and 704 aa, respectively, and have been found to be expressed in almost all tissues in humans ^28,29^. In contrast, isoform *RPGR^ORF15^*, encoding a protein of 1152 aa, has been reported to be retina-specific ^25,28,30,31^. All three major isoforms share the N-terminal part of the protein (regulator of chromosome condensation (RCC)-like domain) ^28,32-34^, which has been associated with terminal part of isoforms RPGR^1-19^ and RPGR^s14/15^ includes an isoprenylation CAAX motif, which is crucial for RPGŔs ciliary localization ^34^.

The physiological roles of isoform *RPGR^1-19^* and isoform *RPGR^ORF15^* in the retina have been comparatively well elucidated ^25,30,31,33,37-40^. This is in contrast to the isoform *RPGR^s14/15^*, which has received limited attention. Notably, isoforms orthologous to the human *RPGR^1-19^* and *RPGR^ORF15^* are present in vertebrate species, including, e.g., Dog, Mice, Zebrafish, and Bovine ^25,30,31,33,37-45^; however, the isoform *RPGR^s14/15^*seems to be expressed only in humans. We previously showed that the human *RPGR* major isoforms build different interaction complexes with other ciliary proteins, indicating distinct roles of the different RPGR isoforms ^29^. The isoform balance seems crucial for normal RPGR functions, further emphasizing their unique properties ^46-48^.

Primary cilia were primarily considered a microtubule-based organelle; however, increasing evidence suggests that also actin is relevant for ciliogenesis and cilia maintenance ^49-51^. Recently, actin molecules, particularly filamentous actin (F-actin), were shown to localize within the cilium, where they play a role in organizing the ciliary assembly and disassembly ^52,53^. Furthermore, pharmacological modulators of actin polymerization, such as Cytochalasin D (CytoD) and Latrunculin A (LAT-A), perturb normal actin turnover and influence the ciliary length and maintenance ^14,53-57^.

Previous studies have shown that actin turnover is involved in regulating the photoreceptor outer segment renewal process ^58-60^. Megaw and colleagues recently found that the loss of the two major murine *Rpgr* isoforms (*Rpgr^1-18^* and *Rpgr^ORF14^)* from a knockout model disturbed the normal actin turnover process and led to the formation of hyperstabilized F-actin stress fibers, which in turn hinders the disc genesis process of photoreceptor rods and cones ^58^. Premature shedding of newly formed outer in the *Rpgr* knockout mouse ^58^. Notably, a 6h short-term pharmacological modulation of actin hyperstabilization using CytoD or a LIM kinase inhibitor partially corrected the phenotype ^58^.

Actin turnover is tightly regulated by Rho-family GTPases, e.g., RhoA, Rac1, and Cdc42, which control F-actin polymerization, depolymerization, severing, and branching ^61,62^. Rho GTPase activation regulates the Rho-associated coiled-coil containing protein kinases 1 and 2 (ROCK1 and 2)-mediated phosphorylation of LIM kinases. LIM kinases than phosphorylate cofilin to regulate actin severing and depolymerization ^63,64^. Concurrently, activated Cdc42 and Rac1 promote actin polymerization and branching via interactions with N-WASP and WAVE, respectively, leading to the formation of branched actin networks via the Arp2/3 complex ^62,65,66^.

Ripasudil is a small compound that is a potent inhibitor of ROCK 1 and ROCK 2 kinases with 50 % inhibitory concentrations (IC50) of 0.051 and 0.019 μmol/L, respectively ^67,68^. Ripasudil treatment has been shown to manipulate actin dynamics both *in vivo* and *in vitro* ^68,69^. Notably, Ripasudil 0.4 % ophthalmic solution was approved in Japan for the treatment of glaucoma in 2014 and is currently being investigated for other diseases such as diabetic retinopathy and Retinopathy of prematurity ^70-74^.

To overcome *RPGR*-related defects, we herein investigated the therapeutic potential of the human-specific *RPGR* isoform *RPGR^s14/15^*as well as the clinically approved Rho/ROCK pathway inhibitor Ripasudil. We evaluated ciliary properties and actin turnover in *RPGR* mutant hTERT-RPE1 cells. Our study increases the understanding of treatment options in *RPGR*-associated RP.

## Results

### Generation of isoform-specific *RPGR* mutant hTERT RPE-1 cell lines using CRISPR-Cas9

To investigate the ciliary functions of *RPGR^1-19^* and *RPGR^s14/15^* isoforms, we generated isoform-specific *RPGR* mutant hTERT-RPE1 cell lines using CRISPR-Cas9 genome editing. To generate a knockout (KO) of all *RPGR* isoforms (RPGR_KO1), we applied a single guide RNA (sgRNA) targeting *RPGR* exon 2 that induced a homozygous 1 bp duplication *RPGR*:c.(114dupA) in exon 2. The mutation leads to a frameshift and is predicted to cause a premature stop codon (p.(His39Thrfs*7)) (Figure 1A and Figure S1A). We compared this cell line to a second knockout cell line (RPGR_KO2, homozygous mutation *RPGR*:c.(122dupC) in exon 2, kindly provided by Prof. Dr. Val C Sheffield and Zhang Qihong, Institute for Vision Research, University of Iowa Carver College of Medicine) ^34^. RPGR_KO1 and RPGR_KO2 show a distinct 1 bp duplication in exon 2 of *RPGR*, but cause a premature stop codon at the same position in *RPGR* (p.(Ser41Serfs*5)) (Figure 1A and Figure S1A).

**Figure 1.**
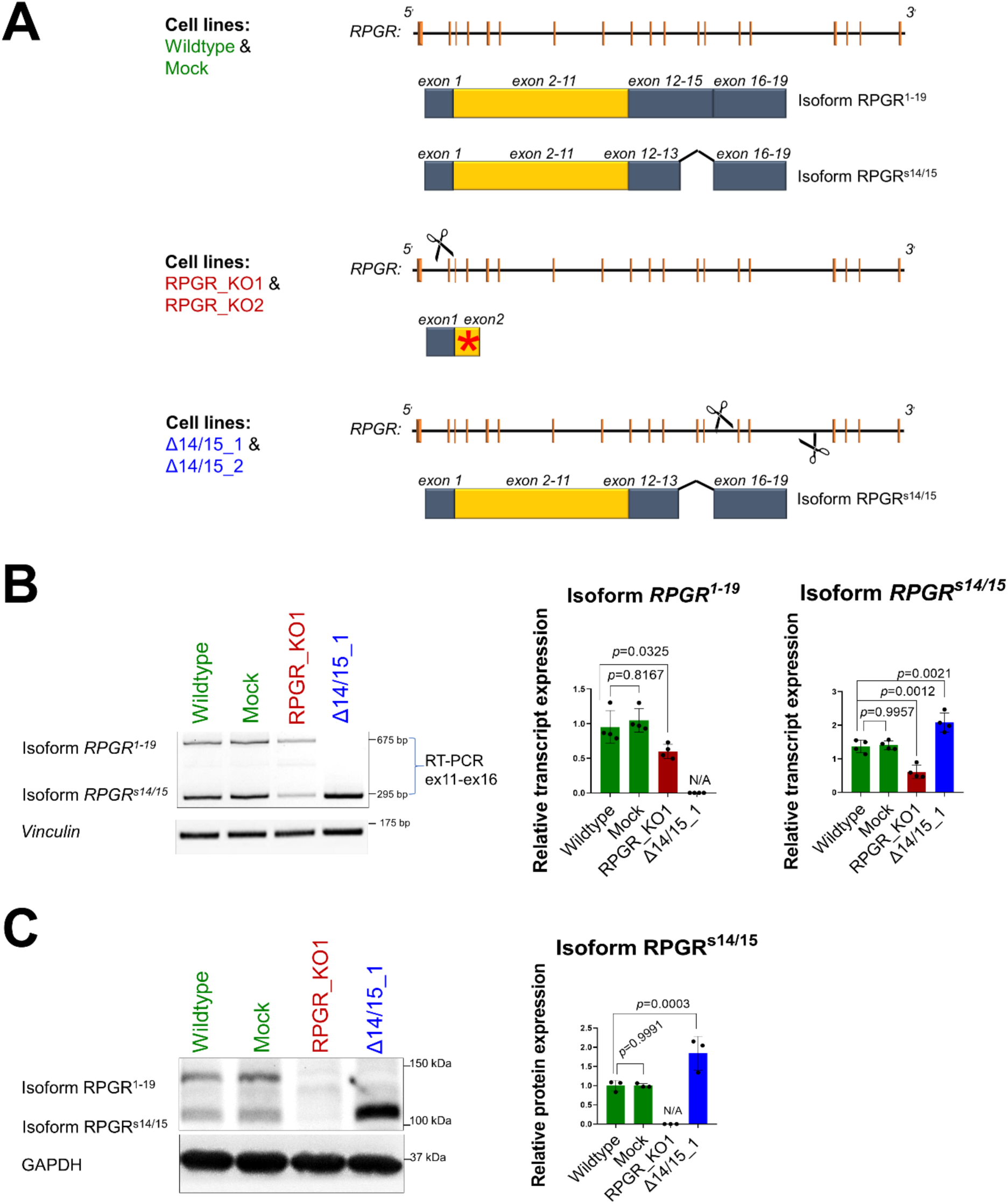
Generation of hTERT RPE1 cell lines with mutations in *RPGR*. **A.** Schematic diagram summarizing the genetic bases of CRISPR-modified RPE1 cell color vertical lines between the 5 ′ and 3 ′ ends of the *RPGR* gene. The reference cell lines (Wildtype and Mock, in green) express both *RPGR* isoforms, *RPGR^1-19^* and *RPGR^s14/15^*. In cell lines RPGR_KO1 and RPGR_KO2 (in red), a CRISPR-mediated mutagenesis using a single sgRNA (scissor) that targeted exon 2 of *RPGR* resulted in a frameshift and premature stop codon (red asterisk). The mutations affected all *RPGR* isoforms and resulted in RPGR knockouts in two independent cell lines. In the lowest panel, the combination of two sgRNAs (scissors) targeted intron 13 and intron 15 which induced mutations in cell lines Δ14/15_1 and Δ14/15_2 (in blue) leading to the exclusive expression of isoform *RPGR^s14/15^*. **B**. RT-PCR analyses of transcripts from RPE1 wildtype, Mock, RPGR_KO1, and Δ14/15_1 cell lines. Primers that are specific to exons 11 and 16 (ex11-ex16) co-amplified isoforms *RPGR^1-19^* and *RPGR^s14/15^*. *Vinculin* served as a loading control. The size of the PCR products is provided in base pairs (bp). Semi-quantitative densitometric measurements of the RT-PCR band intensities relative to *Vinculin* are shown in the middle and right panels. **C.** Western blot analysis of RPGR proteins in mutant and control RPE1 cells. Proteins of isoforms RPGR^1-19^ and RPGR^s14/15^ were detected in control cells (Wildtype and Mock), but not in the RPGR_KO1 cell line. Cell line Δ14/15_1 exclusively expressed RPGR^s14/15^. Sizes of protein markers are provided in kilodaltons (kDa). Semi-quantitative densitometric measurements of RPGR from Western blots are shown in the right panel. Experiments were independently repeated three times (n=3). One-way ANOVA was used to calculate the *p*-values (non-significant: *p* ≥ 0.05; significant: *p* < 0.05).

Two additional cell lines were generated with the aim of exclusively expressing the isoform *RPGR^s14/15^* (cell line Δ14/15_1 and Δ14/15_2) (Figure 1A). We applied a combination of two sgRNAs targeting intron 13 and intron 15. In the first cell line Δ14/15_1, a compound heterozygous deletion of 4222 bp (on allele 1) and 4228 bp (on allele 2) within the *RPGR* genomic region was identified (Figure S1B). In the second cell line Δ14/15_2, a heterozygous deletion of 4221 bp from intron 13 to intron 15 was identified on one allele, while the second allele showed a 32 bp insertion in intron 13 (Figure S1C).

We screened the RPE1 cell lines for possible off-target sequence alterations induced by the CRISPR-Cas9 treatment. We considered one to three mismatches of the sgRNA with the human genome as potential off-target cutting sites (Table S1). We did not detect sequence alterations using Sanger sequencing of potential off-target sites in the genome of our CRISPR-modified cell lines. In summary, our results indicate that the CRISPR-modified RPE1 cell lines show target site-specific sequence alterations in *RPGR*.

To verify the expression of *RPGR* transcripts in the cell lines studied herein, we used RT-PCR and co-amplified *RPGR* isoform transcripts. In RPGR_KO1, we detected the transcripts of isoform *RPGR^1-19^*and *RPGR^s14/15^* (Figure 1B). The transcript amounts were lower than in controls, possibly because of nonsense-mediated decay of the mutated RNA. We verified that the homozygous point mutation in exon 2 is present in transcripts of RPGR_KO1 (Figure S2A). The cell line Δ14/15_1 showed increased levels of *RPGR^s14/15^* in comparison to controls (Figure 1B). Semi-quantitative densitometric measurements of the RT-PCR band intensities confirmed these observations (Figure 1B). We did not detect the *RPGR* transcript in the cell line RPGR_KO2 (Figure S2B). Cell line Δ14/15_2 resembled Δ14/15_1 (Figure S2B).

Next, we assessed the expression of RPGR protein isoforms in all CRISPR-modified cell lines using western blotting. We found that the RPGR protein was undetectable in RPGR_KO1 and RPGR_KO2, supporting that these cell lines do not express any *RPGR* isoform (Figure 1C and Figure S2C). The Δ14/15_1 and Δ14/15_2 cell lines did not show detectable levels of the RPGR^1-19^ protein isoform. Semi-quantitative densitometric measurements of the band intensities displayed a significant increase (approximately 2 times) in the protein isoform RPGR^s14/15^.

We conclude that RPGR_KO1 and RPGR_KO2 do not express *RPGR* isoforms, whereas Δ14/15_1 and Δ14/15_2 exclusively express the *RPGR^s14/15^* isoform.

### Isoform RPGR^s14/15^ locates to primary cilia

We tested the presence of RPGR proteins in primary cilia and compared our CRISPR-modified cell lines (RPGR_KO1, RPGR_KO2, Δ14/15_1, and Δ14/15_2) with controls (Wildtype, Mock). In Wildtype, Mock, as well as in both Δ14/15 cell lines, approximately 85% of cilia were RPGR positive, indicating that the RPGR^s14/15^ isoform localizes to the cilia (Figure 2A and B). In contrast, ciliary RPGR localization was significantly reduced in both RPGR_KO1 and RPGR_KO2 cell lines. Notably, we detected RPGR staining in approximately 5–10% of cilia in both RPGR_KO1 and RPGR_KO2. This likely reflects coincidental non-specific antibody staining within the cilia of these lines.

**Figure 2.**
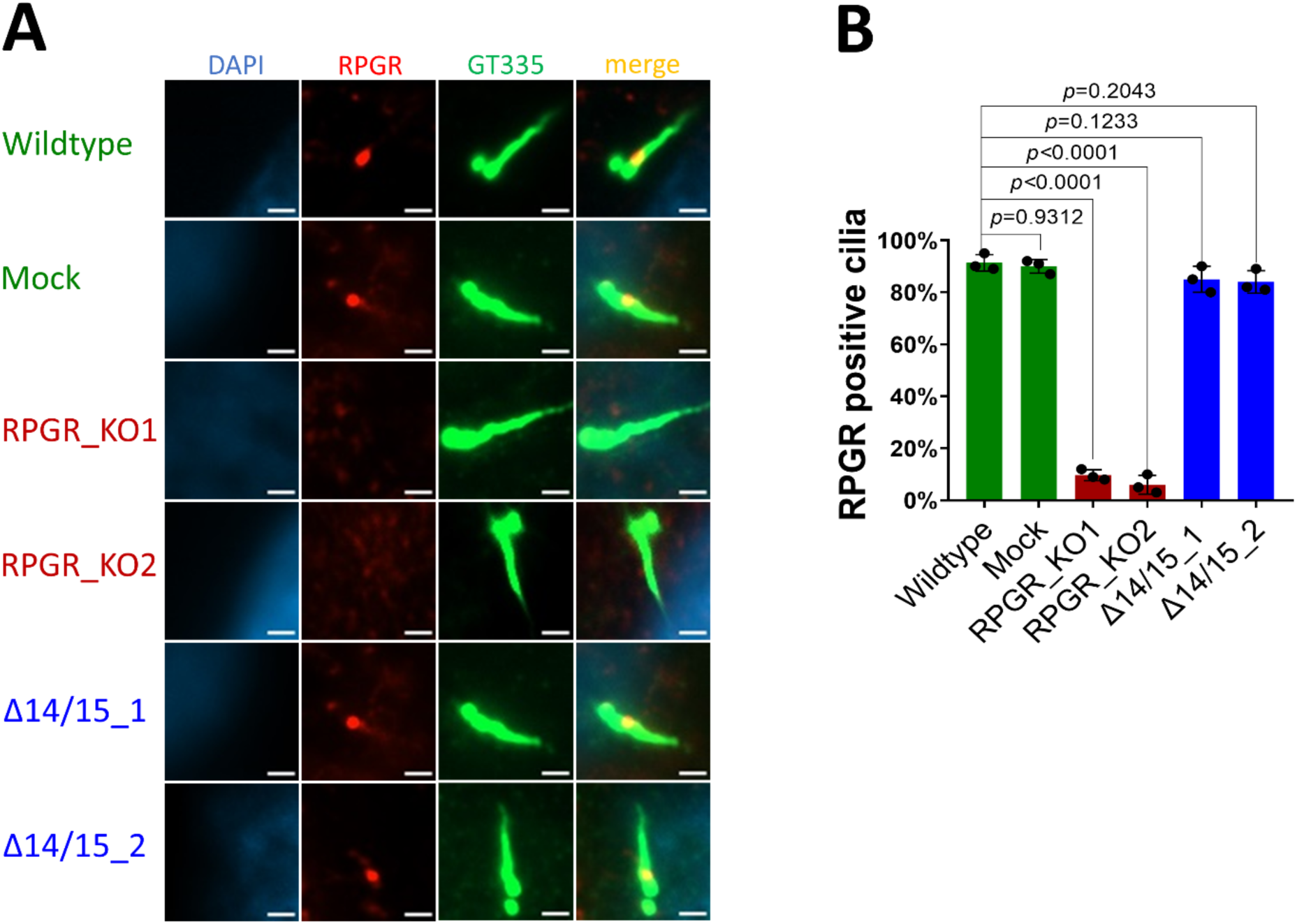
RPGR ciliary localization. **A**. Ciliary localization of RPGR proteins in the cell lines Wildtype, Mock, RPGR_KO1, RPGR_KO2, Δ14/15_1, and Δ14/15_2 analyzed by immunocytochemistry. Cell nuclei were stained with DAPI (blue). RPGR localization was detected using an anti-RPGR antibody (red). The ciliary axoneme was detected using the polyglutamylated tubulin marker GT335 (green). **B**. Quantification of RPGR-positive cilia in percent. We analyzed 50 cilia per experiment and cell line. Experiments were independently repeated three times (n=3). One-way ANOVA was used to calculate the *p*-values (non-significant: *p* ≥ 0.05; significant: *p* < 0.05)—scale bar: 2 μm.

We conclude that the protein of the RPGR^s14/15^ isoform is located in the cilium.

### Isoform RPGR^s14/15^ preserves ciliary length and segmentation

We investigated the relevance of *RPGR* isoforms for ciliary length and segmentation. Cell lines were serum-starved for 48 hours and labeled with two ciliary markers, ARL13B and GT335 (Figure 3A). We measured the length of primary cilia as well as the length of the ciliary proximal segments using the markers ARL13B and GT335, respectively (Figure 3A). We detected significantly increased cilia lengths in *RPGR^s14/15^* isoform (Δ14/15_1 and Δ14/15_2) showed values comparable to controls (Wildtype and Mock) (Figure 3B).

**Figure 3.**
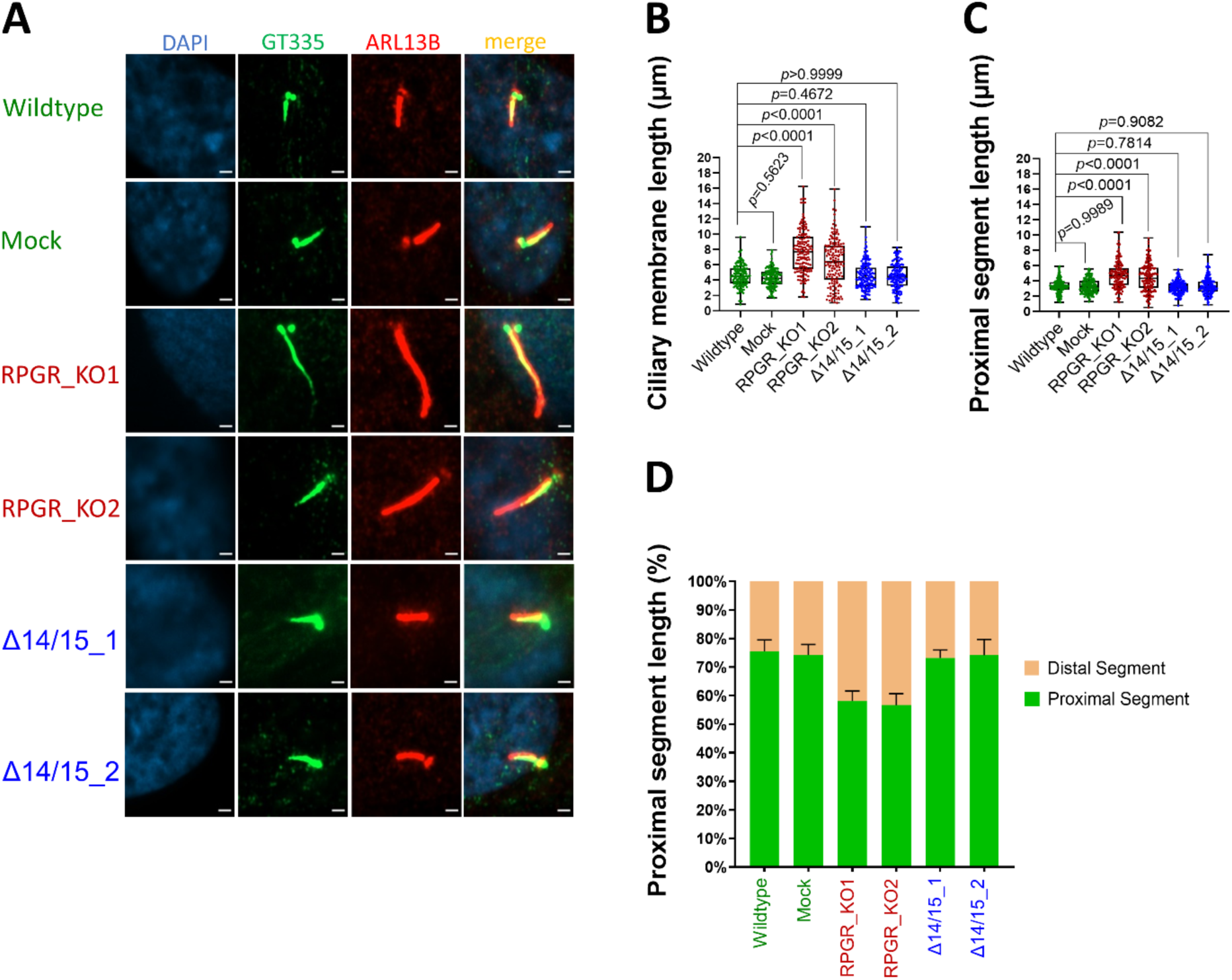
Ciliary segmentation defects in *RPGR* KO mutants. **A**. Immunofluorescence detection of ciliary membrane and ciliary proximal segments. The proximal segment (PS) was stained using polyglutamylated tubulin marker GT335 (green). The ciliary membrane was stained with the marker ARL13B (red). DAPI was used to detect cell nuclei (blue). **B-C.** Quantifications of the length of primary cilia membranes (**B**) and proximal segment length (**C**). **D.** PS length in percent of the cilium was calculated and presented in comparison with the DS. Experiments were independently repeated three times (n=3), and 50 cilia per n were analyzed using Fiji-ImageJ software. One-way ANOVA using GraphPad Prism was used to calculate the *p-*values (non-significant: *p* ≥ 0.05; significant: *p* < 0.05)—scale bar: 2 μm.

Previously, it was reported that ciliary defects influence ciliary segmentation, the ratio of the ciliary proximal segment (PS) and distal segment (DS) ^14^. We investigated whether the *RPGR* isoforms affect ciliary segmentation and measured the PS length (GT335 staining) (Figure 3A and C). We found that the RPGR_KO1 and RPGR_KO2 cell lines, compared to the controls, display significantly elongated PS (Figure 3A and C). We also calculated the PS and DS lengths in percent of the cilium length and observed reduced PS in RPGR_KO1 and RPGR_KO2. This indicated that the ratio of the PS and DS was shifted in cells lacking all *RPGR* isoforms (Figure 3D). Interestingly, cell lines only expressing the isoform *RPGR^s14/15^* (Δ14/15_1 and Δ14/15_2) did not show alterations of PS length (Figure 3C and D) and were comparable to controls.

Our data showed perturbed ciliary length and segmentation ratios in RPGR knockout cells. In contrast, cells exclusively expressing the isoform *RPGR^s14/15^*were indistinguishable from controls. The isoform *RPGR^s14/15^* seems to preserve the ciliary length and segmentation.

### Isoform *RPGR^s14/15^* as well as the Rho/ROCK inhibitor Ripasudil preserve actin turnover

We investigated whether the human-specific isoform *RPGR^s14/15^* contributes to the regulation of actin turnover in the cellular human *RPGR* models studied herein. We used phalloidin staining as a marker for F-actin, and observed that approximately 30% of control cells (Wildtype and Mock) displayed hyper-stabilized F-actin stress fibers (Figure 4A and B), indicating a normal F-actin turnover. In contrast, the KO cell lines (RPGR_KO1 and RPGR_KO2) exhibited a significantly higher percentage of cells, approximately 70% with hyper-stabilized F-actin stress fibers, suggesting an impairment in actin turnover upon loss of all *RPGR* isoforms (Figure 4A and B). Significantly, cell lines exclusively expressing the isoform *RPGR^s14/15^* (Δ14/15_1 and Δ14/15_2) presented with comparable values as controls (Figure 4A and B). These findings suggested that the isoform *RPGR^s14/15^* is capable of preserving normal actin turnover.

**Figure 4.**
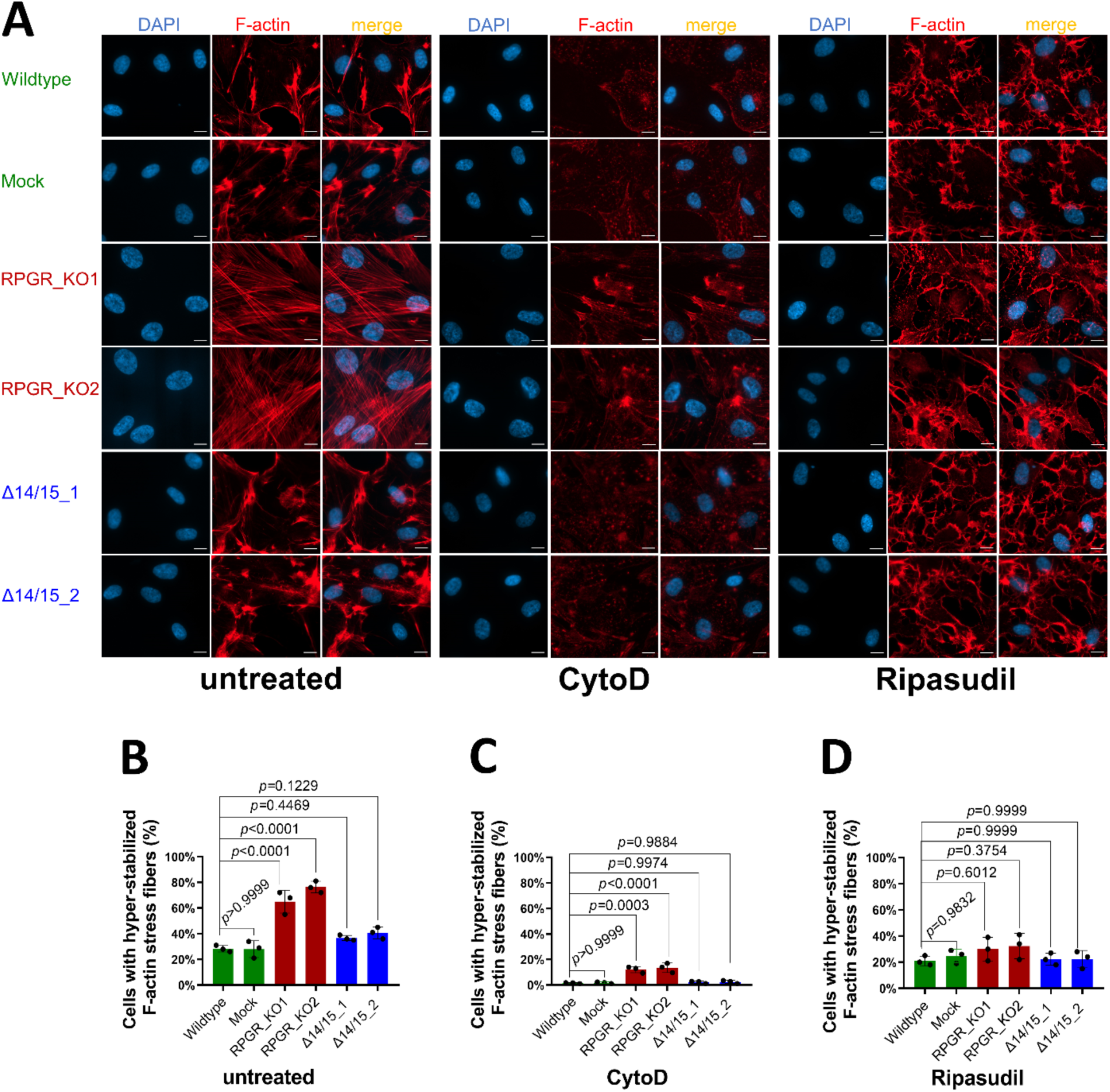
Defective actin turnover is caused by the loss of *RPGR* and can be treated with the Rho/ROCK inhibitor Ripasudil. **A**. Immunofluorescence detection of F-actin staining (red) using phalloidin in untreated, CytoD (200nM) treated, and Ripasudil (30 µM) treated conditions. Control RPE1 cells (Wildtype and Mock) were compared to RPGR knock-out cells (RPGR_KO1 and RPGR_KO2) as well as to cells exclusively expressing the human-specific *RPGR* isoform *RPGR^s14/15^* (Δ14/15_1 and Δ14/15_2). **B**. Quantitative assessment of cells positive for hyper-stabilized F-actin stress fibers in untreated conditions. **C**. Quantitative assessment of cells positive for hyper-stabilized F-actin stress fibers in CytoD-treated conditions. **D**. Quantitative assessment of cells positive for hyper-stabilized F-actin stress fibers in Ripasudil-treated conditions. Cells showing more than five F-actin stress fibers across the nucleus were considered hyper-stabilized F-actin-positive. DAPI was used to detect cell nuclei (blue). Experiments were independently repeated three times (n=3). A minimum of 250 cells were quantified. One-way ANOVA was used to calculate *p-*values (non-significant: *p* ≥ 0.05; significant: *p* < 0.05)—scale bar: 20 μm.

Recently, loss of all *Rpgr* isoforms was associated with hyper-stabilized actin stress fibers in the outer segment of the mouse retina, a process that perturbed the photoreceptor disc genesis ^58^. Short-term inhibition of F-actin polymerization for 6h by intravitreal application of CytoD partially rescued the mouse phenotype ^58^. We evaluated whether the actin modulator CytoD was also capable of correcting the ciliary and F-actin phenotypes in our KO cell lines. We found that the CytoD treatment significantly changed the appearance of hyper-stabilized F-actin stress fibers in our KO cell lines (RPGR_KO1 and RPGR_KO2); however, it disrupted the actin cytoskeleton in all tested cell lines including the two controls (Wildtype and Mock) as well as both *RPGR^s14/15^* expressing cell lines (Δ14/15_1 and Δ14/15_2) (Figure 4A and C). Consequently, we found that the CytoD treatment seemed to disturb the physiological balance of F-actin polymerization. These results suggested significant cellular side effects due to the CytoD treatment in RPE1 cells. Indeed, CytoD directly binds to actin monomers and inhibits their polymerization into filaments, which seemed to have caused CytoD-mediated actin polymerization dysbalances.

We evaluated an alternative treatment option for F-actin polymerization defects and tested the Rho/ROCK pathway inhibitor Ripasudil. Of note, Ripasudil is a clinically approved drug for Glaucoma ^72^. We found that the Ripasudil treatment significantly reduced the hyperstabilized F-actin stress fibers and restored the normal actin turnover in KO cell lines (RPGR_KO1 and RPGR_KO2) (Figure 4A and D). In contrast to the CytoD treatment, we detected that Ripasudil does not disrupt the actin cytoskeleton in controls (Wildtype and Mock) and in *RPGR^s14/15^*expressing cell lines (Δ14/15_1 and Δ14/15_2) (Figure 4A and D). While CytoD treatments led to significant F-actin-related side effects, Ripasudil seemed to support a well-balanced actin cytoskeleton turnover.

### Rho/ROCK pathway inhibitor Ripasudil rescued ciliary defects associated with the loss of *RPGR*

We tested whether F-actin modulations, either using CytoD or Ripasudil treatments, can correct ciliary defects in RPGR KO cell lines. We found that CytoD treatments, although influencing the F-actin polymerization, did not correct ciliary length and segmentation alterations in KO cells (Figure 5, also compare to Figures 3 and 4). Significantly, the treatment with CytoD elongated the cilia of controls and Δ14/15 cell lines (Figure 5A), phenocopying the defects found in KO cell lines; these observations are consistent with previous findings ^14,55^. The cilia elongation was observed in both the ciliary membrane length and the length of the proximal segment (Figure 5B and C). Additionally, we showed that the CytoD treatment caused a proportional alteration in the proximal segment length percentage of the cilium across all cell types (Figure 5D). These findings confirmed that F-actin alterations impact ciliary properties and additionally indicated strong ciliary side effects of CytoD treatments.

**Figure 5.**
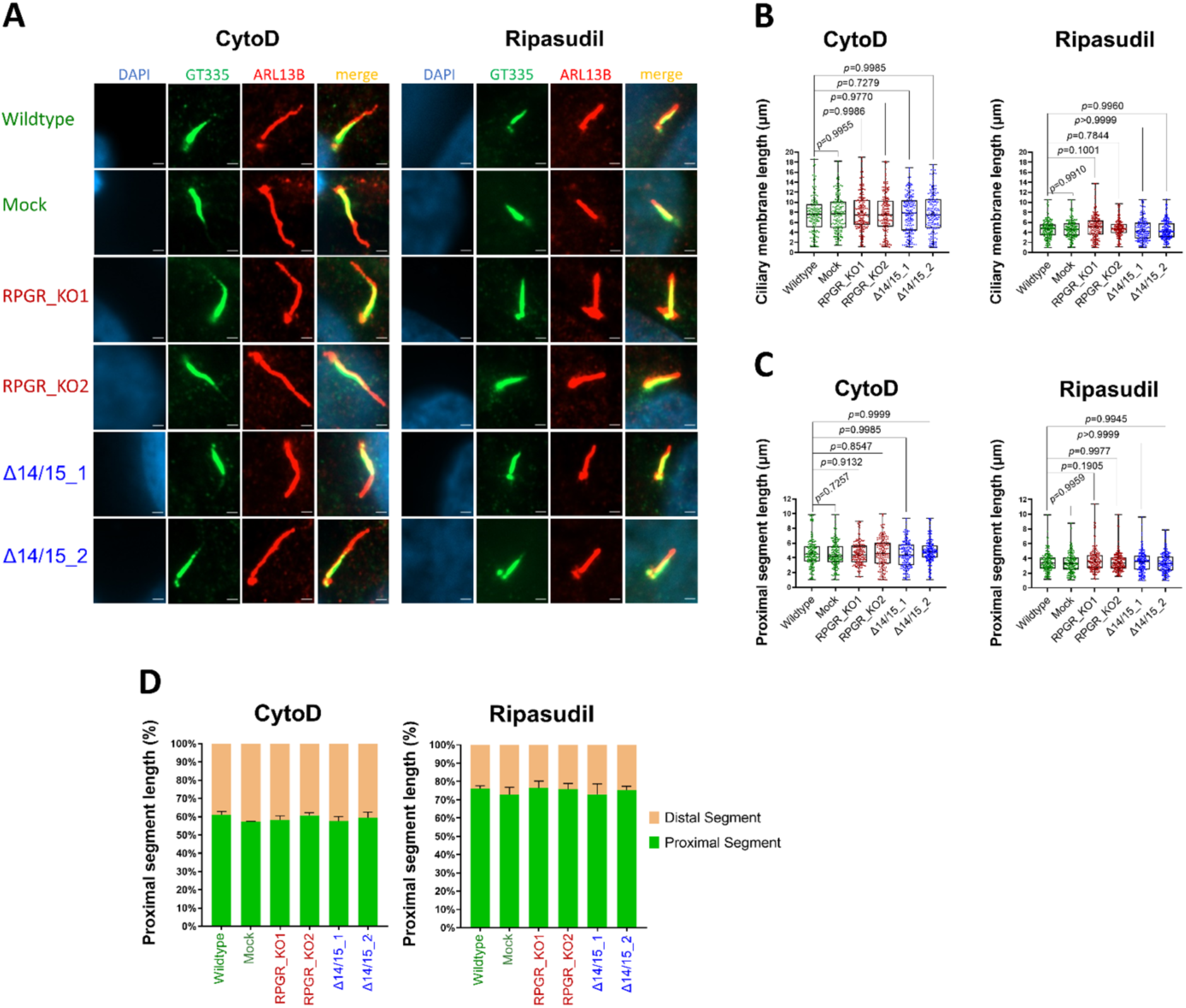
Ripasudil treatment corrects *RPGR*-associated ciliary defects. **A**. Immunofluorescence detection of ciliary membrane and ciliary proximal segments after CytoD (200 nM) or Ripasudil (30 µM) treatments. The proximal segment (PS) was stained using the polyglutamylated tubulin marker GT335 (green). The ciliary membrane was stained with the marker ARL13B (red). DAPI was used to stain cell nuclei (blue). Controls cell lines (Wildtype and Mock) were compared with *RPGR* knockout cells (RPGR_KO1 and RPGR_KO2), as well as cells exclusively expressing the human-specific *RPGR* isoform *RPGR^s14/15^*(Δ14/15_1 and Δ14/15_2). **B-C.** Quantifications of primary cilia membrane lengths **(B)** and proximal segment lengths **(C)** upon CytoD and Ripasudil treatments**. D**. PS length in percent of the cilium length following the treatments with CytoD and Ripasudil. Experiments were independently repeated three times (n=3). More than 160 cilia were analyzed in total. One-way ANOVA using GraphPad Prism was used to calculate the *p-*values (non-significant: *p* ≥ 0.05; significant: *p* < 0.05)—scale bar: 2 μm.

Rho/ROCK pathway inhibitors are well known to influence actin polymerization. We hypothesized that Rho/ROCK inhibition is capable of re-establishing the balance of actin turnover. Treatment of our cell lines with Ripasudil restored the ciliary membrane length as well as the proximal segment length in the cell lines lacking all *RPGR* isoforms (RPGR_KO1 and RPGR_KO2) (Figure 5A-D). In contrast to the CytoD treatments, we detected that Ripasudil did not alter ciliary lengths and segmentation in controls (Wildtype and Mock) and in *RPGR^s14/15^* expressing cell lines (Δ14/15_1 and Δ14/15_2) (Figure 5A-D).

In conclusion, the Rho/ROCK inhibitor Ripasudil presented as a promising therapeutic option to treat *RPGR*-mediated ciliary and actin defects.

## Discussion

The primary cilium orchestrates cellular signaling during development and tissue homeostasis ^8,75^. Defects in actin turnover that lead to ciliary length alterations were previously associated with several ciliopathies, underscoring the importance of an actin-mediated ciliary structure for tissue maintenance ^53,57,58^. *RPGR* is known to influence ciliary structure and the actin cytoskeleton ^27,47,57,58,76,77^; however, the roles of specific *RPGR* isoforms in ciliary structure and actin cytoskeleton remain unclear.

Orthologous isoforms of human *RPGR^1-19^* and *RPGR^ORF15^*have been identified in several species, including mice, zebrafish, and dogs ^25,31,38,78,79^. We have previously shown that human tissues express an additional *RPGR* isoform, *RPGR^s14/15^*, which has not been detected in other species so far ^28,46^. The relevance of these three major *RPGR* isoforms, particularly isoform *RPGR^s14/15^*, is not well understood. Therefore, we investigated the role of the *RPGR^s14/15^* isoform for primary cilia structure and actin cytoskeleton.

Actin turnover plays an important role in maintaining the ciliary length and structure, and F-actin inhibition via CytoD leads to elongated ciliary length ^14,50,55^. We observed that *RPGR* KO cells exhibit elongated cilia along with elongated proximal segment length. Thus, loss of all *RPGR* isoforms perturbed ciliary segmentations (alterations in lengths of PS and DS). Such structural changes in the primary cilia have been correlated with actin or tubulin polymerization/depolymerization defects ^14,50,52^. Interestingly, cells exclusively expressing the isoform *RPGR^s14/15^* did not show any such alterations in ciliary length or segmentation, which suggested that the *RPGR^s14/15^*isoform is likely relevant for stabilizing ciliary architecture, maybe via the regulation of actin dynamics. Additionally, we also show that CytoD treatment impacts ciliary length for ciliary stability ^14,55,80^. The loss of all *RPGR* isoforms (RPGR_KO1 and RPGR_KO2) disrupted the normal actin turnover, a critical process in maintaining cellular and ciliary properties. Vice versa, actin-altering small compounds (like CytoD) can induce ciliary architectural defects. Exclusive expressions of the isoform *RPGR^s14/15^*seem to maintain both the normal cellular actin turnover and the ciliary architecture, suggesting its potential as a therapeutic option to treat *RPGR*-associated RP. Integrating the previously published data with our findings, we speculate that an imbalance between the *RPGR* isoforms causes alterations in actin-mediated axoneme stability. We previously described that isoform *RPGR^s14/15^*builds different interaction complexes with known *RPGR* binding partners, a finding that might be relevant in this context ^46^. Additionally, our data show that isoform *RPGR^s14/15^* localizes to primary cilia.

In line with other groups, we observed that the loss of all *RPGR* isoforms leads to hyper-stabilized F-actin stress fibers ^57,58,76,77^. In *RPGR* knock-out mouse retinae, Megaw et al. recently found that actin stress fibers impact the photoreceptor disc genesis ^58^. They showed that a short-term treatment with CytoD or with a LIM kinase inhibitor ameliorated the hyper-stabilized actin stress fibers in the connecting cilium of photoreceptors. Nevertheless, these treatments caused an overgrowth of the basal discs in the *Rpgr* KO murine model ^58^. In agreement with these findings, our analyses in *RPGR* KO cell lines confirmed that CytoD treatment can dissolve hyper-stabilized F-actin stress fibers; however, it severely impacted the actin cytoskeleton balance and consequently changed the ciliary architecture, also in control cells. Our data suggested that F-actin modulation with CytoD treatment has substantial limitations as a therapeutic option to treat *RPGR*-associated defects.

We found that the inhibitor of the Rho/ROCK pathway Ripasudil, a clinically-approved drug for the treatment of Glaucoma, mediated benefits to the normal actin turnover and rescued the ciliary defects associated with *RPGR* loss. Interestingly, recent findings from other diseases or ciliopathies supported that ciliary and actin-based defects benefit from Rho/ROCK pathway inhibition ^81,82^. To the best of our knowledge, our data suggested for the first time that Ripasudil ameliorates *RPGR*-associated actin and ciliary defects without obvious side effects. As Ripasudil is commercially available, primarily approved for the treatment of glaucoma and ocular hypertension ^67,72,73^, the existing formulation might offer an avenue for repurposing ripasudil to target actin-mediated retinal defects, e.g. in *RPGR*-associated RP.

Beyond cytoskeletal modulation, ripasudil exhibits anti-inflammatory properties in the retina ^70,83-85^. This dual action—correcting cytoskeletal anomalies and reducing inflammation—positions ripasudil as a promising candidate for treating retinal diseases such as *RPGR*-associated RP. While topical application of ripasudil has shown promising outcomes for other diseases, the potential and efficacy of Ripasudil in treating *RPGR*-associated retinal conditions are yet to be understood in detail. Notably, studies have indicated that following topical administration in rabbits, ripasudil achieves measurable concentrations in the retina and choroid ^86^, suggesting its capacity to traverse ocular barriers. We propose that future research should elucidate Ripasudiĺs efficacies in reverting/ameliorating *RPGR*-associated retinal defects as well as optimizing administration routes.

To the best of our knowledge, we are the first to demonstrate that the isoform *RPGR^s14/15^* is capable of maintaining the actin dynamics and ciliary properties. The exact mechanism by which this isoform influences actin dynamics is not yet clear and requires further investigation. However, we can speculate that isoform *RPGR^s14/15^* may have direct/indirect interactions with key actin regulators, connecting cilia-associated proteins and/or ciliary signaling pathways. *RPGR* was reported to interact with gelsolin and cofilin to regulate the actin dynamics ^58,76^. Indeed, our observations support that actin remodeling is correlated to ciliary phenotypes and leads to impacts on ciliary architecture, also in *RPGR* knock-out cells. The isoform *RPGR^s14/15^* thus appears to be relevant to stabilize ciliary architecture.

In conclusion, our investigations established a novel physiological role of *RPGR-* isoform *RPGR^s14/15^* in maintaining the ciliary architecture and actin dynamics. Furthermore, we show that the Rho/ROCK pathway inhibitor Ripasudil, a clinically approved small molecule, can rescue or ameliorate the *RPGR*-associated actin and ciliary defects. The improved understanding of *RPGR*-associated molecular and cellular defects provides the foundation for therapeutic developments.

## Materials and methods

### Cell culture and treatments

hTERT-immortalized retinal pigment epithelial cells (hTERT-RPE1) were kindly provided by Prof. Dr. Floris Foijer at the University Medical Center Groningen. hTERT-RPE1 were cultured in Dulbecco’s modified Eagle medium (DMEM, high glucose, 4.5g/L glucose, Biowest, Nuaille, France) or DMEM (4.5g/L glucose, PAN Biotech, Aidenbach, Germany) supplemented with 10% fetal bovine serum (Sigma-Aldrich, Merck), 1.3% L-glutamine (Biowest), and 1.3% Pen/Strep (Biowest) and incubated at 37 °C and 5% CO_2_. Cells were regularly maintained and passaged using trypsin (Biowest) in a 1:10 ratio twice a week.

We seeded 2×105 hTERT RPE1 cells for actin detection and 4×105 hTERT RPE1 cells for cilia measurements. Before each assay, cells underwent serum starvation in DMEM medium supplemented with 2% L-glutamine (Biowest) and 1% antibiotic (Biowest) for 48 hours. Subsequently, for CytoD treatments, cells received starvation medium containing 200nM CytoD (C2618-200UL, Sigma-Aldrich, Germany) dissolved in DMSO (Carl Roth, Karlsruhe, Germany) for 3 hours. For Ripasudil treatments, cells were incubated with starvation medium containing 30μM Ripasudil (K-115 hydrochloride dihydrate 5mg, Catalog No.S7995, Selleck Biotechnology GmbH, Cologne, Germany) dissolved in DMSO (Carl Roth, Karlsruhe, Germany) for 1 hour. Controls received only DMSO in the same concentration as the treated samples.

### Generation of RPGR mutant clones using CRISPR-Cas9 system

To generate the *RPGR* mutant cell lines, 0.5 million hTERT-RPE1 cells were seeded in a 6-well plate overnight. The expression vector KA2601 eSpCas9-G2P was obtained from Addgene (#124203, Addgene, Watertown, MA, USA). This vector contains three BbsI (New England Biolabs) cutting sites, which complicates the cloning of sgRNAs due to unintended cleavage. Hence, the plasmid modification was done by removing the additional BbsI site using NruI (NEB) and StuI (NEB) enzymes. To create the KO of all *RPGR* isoforms (designated as RPGR_KO1), a single guide RNA (sgRNA) targeting exon 2 (5′-CTCCACATGAAAGATGTACA-3′) was designed using CHOPCHOP (https://chopchop.cbu.uib.no/) and cloned into the KA2601 eSpCas9-G2P vector using BbsI (NEB). 500,000 hTERT-RPE1 cells were counted and seeded in a 6-well plate and incubated overnight. The transfection of the sgRNA containing plasmid (4 μg) was done using FuGeneHD (Promega) according to the manufacturer’s protocols (3:1 ratio (3μl of FuGeneHD into 1μg of plasmid DNA). After 48 h of transfection, cells were subjected to fluorescence-activated cell sorting (FACS) (S3e cell sorter, BioRad, Hercules, CA, USA) to select COPGFP-positive cells. Following 4-5 weeks of culturing the single-cell sorted hTERT-RPE1 cells, genomic DNA extraction was performed using the Puregene® Blood Core Kit B (Qiagen, Hilden, Germany) as per the manufacturer’s instructions. Subsequently, PCR assays using HotFire Tag Polymerase (Solis Biodyne, Tartu, Estonia) were conducted to screen for deletions of a larger fragment. Purification of the PCR amplicons was achieved using Exo-SAP (New England Biolabs) before subjecting them to Sanger sequencing. The sequencing reactions were performed using the BigDye Terminator v3.1 Cycle Sequencing Kit on an ABI Prism 3130xl Genetic Analyzer (Applied Biosystems). SnapGene software (GSL Biotech LLC) was employed to compare the sequencing profile of each clone with the *RPGR* reference sequence NM_000328.3. The RPGR_KO2 cell line was a gift from Dr. Qihong Zhang and Prof. Dr. Val C. Sheffield’s lab. The cell lines exclusively expressing the single isoform *RPGR^s14/15^* (referred to as Δ14/15_1 and Δ14/15_2) were generated by employing a combination of two sgRNAs targeting intron13 and intron15 CAGTACATTTGGTTAGTTAG-3′)), following our published protocol ^46^, except using FuGeneHD (Promega) instead of polyethyleneimine (PEI) transfection reagent. Finally, potential off-target sites associated with each sgRNA were identified through the CHOPCHOP web tool (https://chopchop.cbu.uib.no/) and screened by PCR assays followed by Sanger sequencing. The details of screened off-target sites and primers are provided in Table S1 and Table S2.

### Western blotting

One million hTERT-RPE1 cells were seeded in a T-75 culture flask and cultured at 37°C, with 5% CO2. Before lysis, cells were serum-starved in DMEM medium supplemented with 2% L-glutamine (Biowest) and 1% antibiotic (Biowest) for 24 hours. Lysis was performed on ice using radioimmunoprecipitation assay (RIPA) buffer (1% NP40, 0.1% sodium dodecyl sulphate (SDS), 0.5% sodium deoxycholate in PBS) supplemented with protease inhibitor cocktail and phosphatase inhibitor cocktails 2 and 3 (Sigma-Aldrich). Afterwards, the lysates were incubated on ice for 20 minutes, followed by centrifugation at 3000g for 5 minutes at 4°C. After centrifugation, the supernatant containing proteins was collected, and the concentration was determined using Quick Start Bradford 1× Dye Reagent (Bio-Rad Laboratories, Munich, Germany). Protein concentrations of all samples were adjusted to the same value, and lysates were mixed with sample buffer (10% glycerol, 1% beta-mercaptoethanol, 1.7% SDS, 6.25% 1 M Tris (pH 6.8), 3.34% bromophenol blue) and heated for 5 minutes at 95 °C. Subsequently, 60 µg of each sample was loaded onto 10% SDS-polyacrylamide gels and separated by electrophoresis. Protein transfer onto Polyvinylidene difluoride (PVDF) membranes (0.45 µm pore size; Millipore, Burlington, VT, USA) was achieved by wet blotting at 45 V for 2 hours. The membrane was then blocked in 5% bovine serum albumin in TBST for 30 minutes and incubated overnight with primary antibodies (rabbit polyclonal RPGR antibody (1:500; HPA001593, Sigma-Aldrichand, and mouse monoclonal GAPDH antibody (1:1000; 0508007983, Chemicon International, Temecula, CA, USA)) at 4 °C. After incubation with horse radish peroxidase (HRP)-linked secondary antibodies (Novus Biological, NBP2-30348H and NB7539, both 1:4000) at room temperature for 2 hours and with enhanced chemiluminescence solution (ECL, Thermo Fisher Scientific), protein bands were visualized using an Intas Ecl Chemocam Imager.

### RNA isolation and transcript expression analysis

Total RNA extraction from cultured hTERT-RPE1 cells was performed using the NucleoSpin RNA kit (Macherey-Nagel, Düren, Germany) following the manufacturer’s instructions. Using 500 ng total RNA in each sample, cDNA was reverse transcribed using random primers and SuperScript III Reverse Transcriptase (Invitrogen, Carlsbad, CA, USA) according to the manufacturer’s recommendations. RT-PCRs were conducted to confirm the presence of *RPGR* transcripts, utilizing HotFire Tag Polymerase (Solis Biodyne, Tartu, Estonia). Subsequently, all RT-PCR products were assessed on a 2% agarose gel and subjected to sequence verification via Sanger sequencing. Primer sequences are provided in Table S2.

### Immunocytochemistry and fluorescence microscopy

For immunocytochemical experiments to investigate ciliary properties, 4×10^5^ hTERT RPE1 cells were seeded in DMEM (Biowest). For actin stainings, 2×10^5^ were seeded in DMEM (PAN Biotech). Cells were either fixed in cold methanol (Sigma-Aldrich, Merck) at −20°C for 5 min (cilia staining) or in 4% PFA (Carl Roth) at room temperature for 10 min (actin staining). After fixation, cells were washed three times with PBS-Tx (1× Phosphate Buffered Saline (Chemsolute) containing 0.1% Triton X-100 incubated for 30 min at 80 °C in 0.1 M Tris-HCl (pH 9) after the fixation with cold methanol at −20°C for 5 min. Cells were washed again twice with the PBS-Tx. Blocking was performed by treating cells with PBS-Tx containing 3% BSA (fraction V; Carl Roth) for 30 min at room temperature in a wet chamber. Blocked samples were then incubated with diluted primary antibodies (cilia staining) overnight at 4 °C, or with Alexa Fluor™ 647 Phalloidin (Thermo Fischer Scientific) for 1h at room temperature in a dark, wet chamber. Afterward, coverslips were washed with PBS-Tx and were incubated with secondary antibodies (not relevant for actin staining) for 1 h at room temperature in a dark wet chamber. Afterward, samples were washed again with PBS-Tx. Finally, the cells were mounted with Fluoromount containing 4′, 6-diamidino-2-phenylindole (DAPI) (SouthernBiotech, Birmingham, AL, USA). All antibodies and Phalloidin were diluted in the blocking solution. Images were acquired using a fluorescence microscope (Axiophot; Carl Zeiss) equipped with an AxioCam camera and Axiovision software. Ciliary axoneme length was measured using Fiji-ImageJ software (ImageJ, https://fiji.sc/). To quantify cells that are positive for hyper-stabilized F-actin thin filaments, we counted the cells showing more than five F-actin filaments at/over the nucleus (stained by DAPI). The list of all antibodies (Primary and secondary) is provided in Table S3.

### Statistics

The data are presented as mean ± SD unless otherwise indicated. Statistical analysis was performed using one-way ANOVA for all data sets, followed by Tukey’s multiple comparison test using GraphPad Prism software (version 9.4.1, www.graphpad.com). Differences between groups were considered nonsignificant if *p* ≥ 0.05, and significant if *p* < 0.05.

## Acknowledgments

We are grateful to Prof. Dr. Floris Foijer from the University Medical Center Groningen and his PhD student, Christy Hong, for generously providing the hTERT-RPE1 cells. We also thank Dr. Qihong Zhang and Prof. Dr. Val C. Sheffield from the Department of Ophthalmology and Visual Sciences, University of Iowa, USA, for providing the knock-out cell line of *RPGR* (RPGR_KO2). Additionally, we express our appreciation to Prof. Dr. Karl-W. Koch from the Department of Biochemistry, School of Medicine and Health Sciences, University of Oldenburg, for his insightful discussions and valuable contributions.

## Author contributions

Conceptualization, J.N.; experimental design, MU.; Methodology, M.U., J.M., D.K., C.J., P.A., J.M.B., C.L.K.; Validation, M.U.; Investigation, M.U., D.K., J.M.B., C.L.K.; Statistical analysis, M.U., C.J.; Resources, J.N.; Data Curation, M.U., J.N.; Writing— Original Draft Preparation, M.U.; Writing—Review & Editing, M.U., C.J., P.A., J.N.; Supervision, J.N.; Project Administration, J.N.; Funding Acquisition, J.N. All authors have read and agreed to the published version of the manuscript.

