## Supplementary data for "The human-specific *RPGR* isoform *RPGR^s14/15^* and the clinically approved Rho/ROCK inhibitor Ripasudil represent therapeutic options to address *RPGR*-associated defects"

**Title**

**Affiliations**

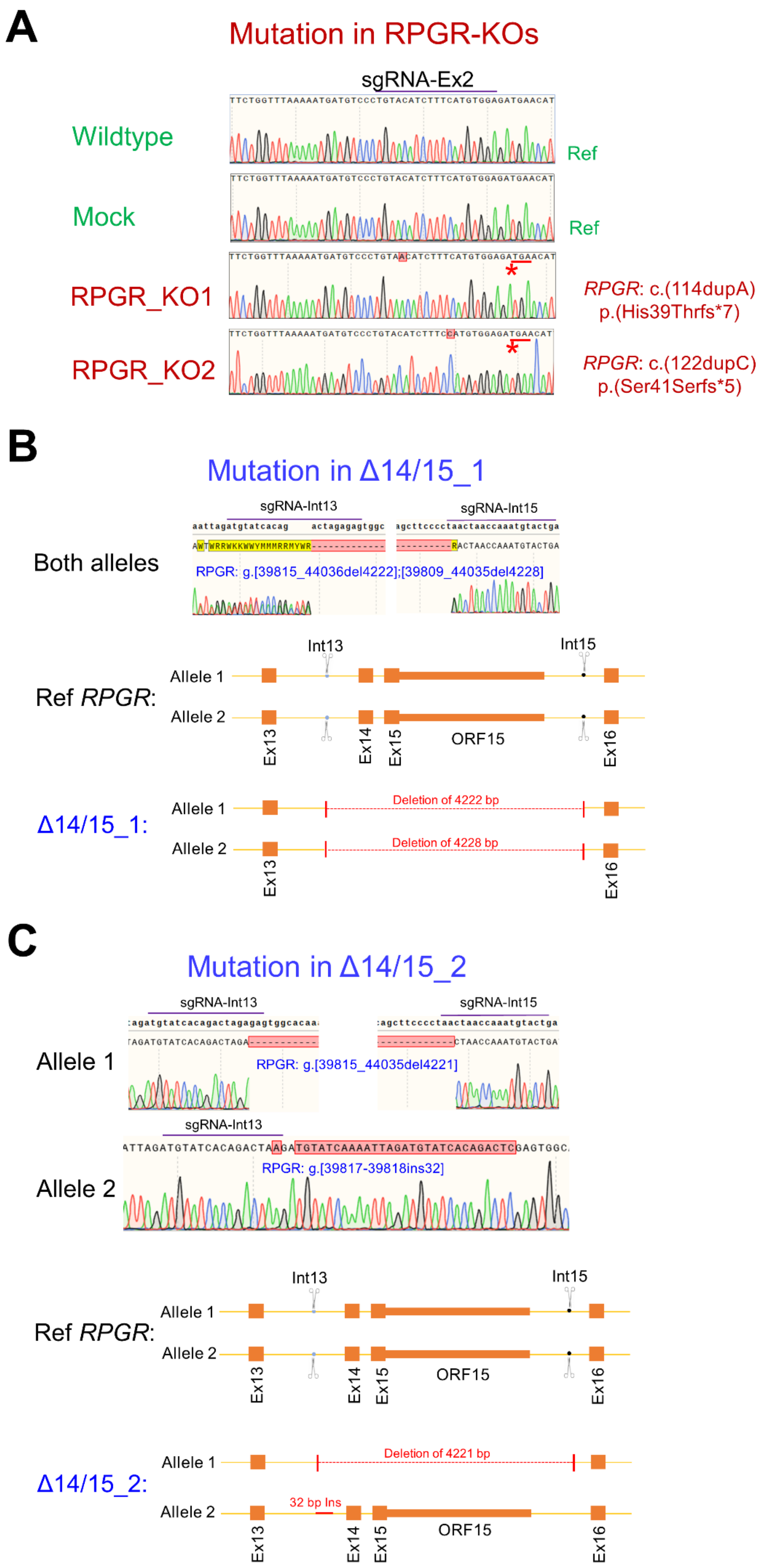
A
Mutation in RPGR-KOs
sgRNA-Ex2
Wildtype
Ref
Mock
Ref
RPGR_KO1
RPGR: c.(114dupA) p.(His39Thrfs*7)
RPGR_KO2
RPGR: c.(122dupC) p.(Ser41Serfs*5)
B
Mutation in Δ14/15_1
sgRNA-Int13
sgRNA-Int15
Both alleles
RPGR: g.[39815_44036del4222];[39809_44035del4228]
Ref RPGR:
Allele 1
Allele 2
Int13
Int15
Ex13
Ex14
Ex15
ORF15
Ex16
Δ14/15_1:
Allele 1
Deletion of 4222 bp
Allele 2
Deletion of 4228 bp
Ex13
Ex16
C
Mutation in Δ14/15_2
sgRNA-Int13
sgRNA-Int15
Allele 1
RPGR: g.[39815_44035del4221]
sgRNA-Int13
Allele 2
RPGR: g.[39817-39818ins32]
Ref RPGR:
Allele 1
Allele 2
Int13
Int15
Ex13
Ex14
Ex15
ORF15
Ex16
Δ14/15_2:
Allele 1
Deletion of 4221 bp
Allele 2
32 bp Ins
Ex13
Ex14
Ex15
ORF15
Ex16

**Figure S1. Genetic characterization of *RPGR* mutant cells. A.** In RPGR_KO1, the guide RNA sgRNA-Ex2 resulted in a homozygous duplication of 1bp in exon 2 (c.114dupA). In RPGR_KO2, the same guide RNA sgRNA-Ex2 induced a different homozygous duplication of 1 bp in exon 2 (c.122dupC). RPGR_KO1 and RPGR_KO2 frame shifts result in the same premature stop codon (red bar and asterisk). **B.** The combined treatment with two sgRNAs (sgRNA-Int13 targeting intron 13 and sgRNA-Int15 targeting intron 15) resulted into deletions of 4222bp (first allele) and 4228bp (second allele) in cell line Δ14/15_1. The lower panel shows schematic drawings of the CRISPR-induced genetic rearrangements. **C.** The combination of the two sgRNAs sgRNA-Int13 and sgRNA-Int15 resulted in the deletion of 4221 bp from the first allele and a 32 bp insertion in the second allele of cell line Δ14/15_2. Schematic presentation of genetic rearrangements in cell line Δ14/15_2 is shown in the lower panel. Ref *RPGR* indicates the reference sequence before CRISPR modification.

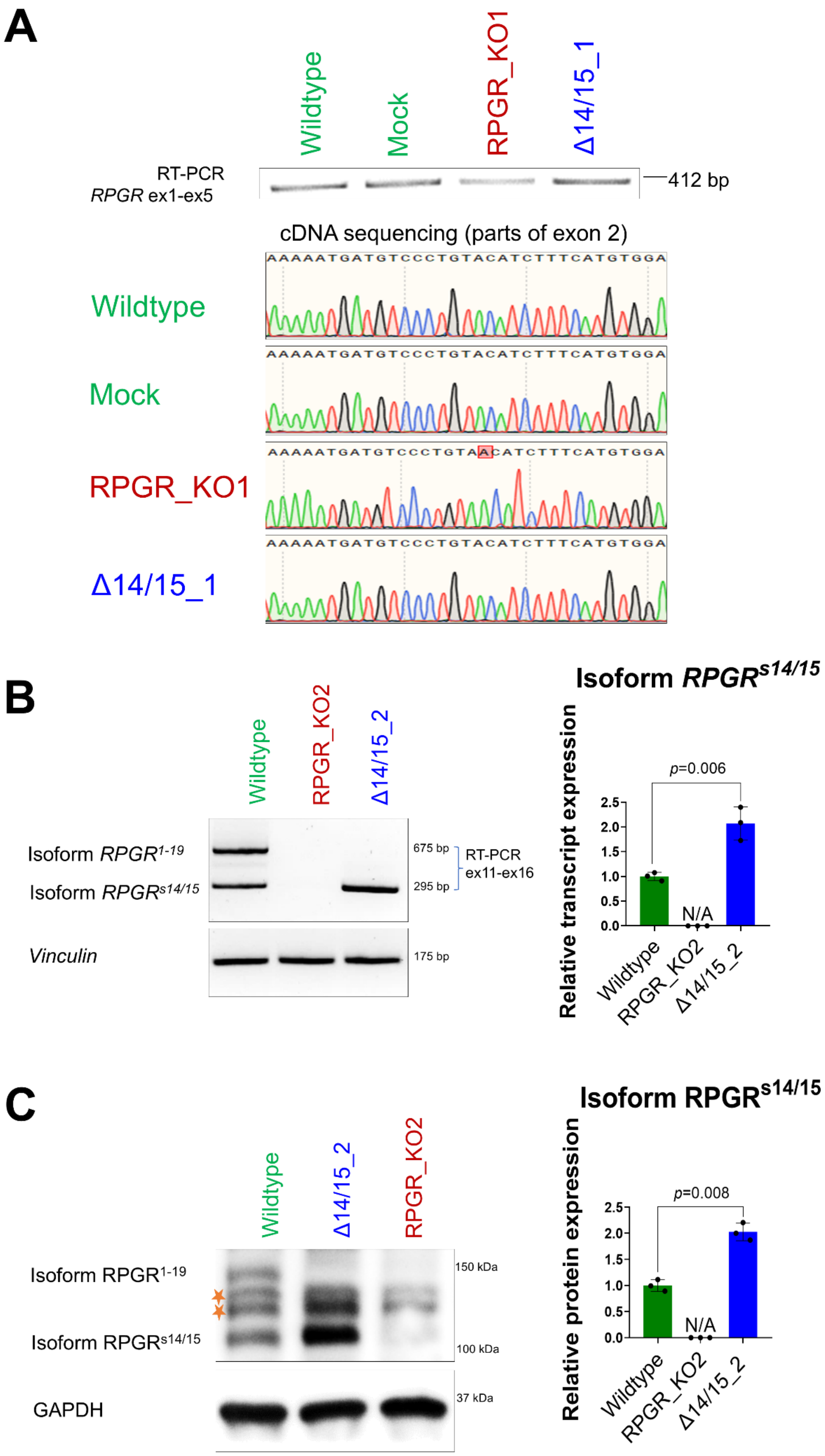

A
Wildtype
Mock
RPGR_KO1
Δ14/15_1
RT-PCR
RPGR ex1-ex5
412 bp
cDNA sequencing (parts of exon 2)
AAAAATGATGTCCCTGTACATCTTTCATGTGGA
Wildtype
AAAAATGATGTCCCTGTACATCTTTCATGTGGA
Mock
AAAAATGATGTCCCTGTAACATCTTTCATGTGGA
RPGR_KO1
AAAAATGATGTCCCTGTACATCTTTCATGTGGA
Δ14/15_1
B
Wildtype
RPGR_KO2
Δ14/15_2
Isoform RPGR1-19
Isoform RPGRs14/15
Vinculin
675 bp
295 bp
175 bp
RT-PCR
ex11-ex16
Isoform RPGRs14/15
Relative transcript expression
p=0.006
2.5
2.0
1.5
1.0
0.5
0.0
N/A
Wildtype
RPGR_KO2
Δ14/15_2
C
Wildtype
Δ14/15_2
RPGR_KO2
Isoform RPGR1-19
Isoform RPGRs14/15
GAPDH
150 kDa
100 kDa
37 kDa
Isoform RPGRs14/15
Relative protein expression
p=0.008
2.5
2.0
1.5
1.0
0.5
0.0
N/A
Wildtype
RPGR_KO2
Δ14/15_2

**Figure S2. Transcript and protein characterization of *RPGR* mutant cell lines. A.** RT-PCR amplification of *RPGR* transcripts from exon 1 to exon 5 from Wildtype, Mock, RPGR_KO1, and Δ14/15_1 cell lines. Sanger sequencing of these *RPGR* transcripts showed reference sequences in Wildtype, Mock, and Δ14/15_1 cell lines. The homozygous sequence alteration c.(114-115insA) in RPGR_KO1 was confirmed. **B.** Isoform-specific RT-PCR assay using primers that bind to *RPGR* exons 11 and 16. Co-amplification of isoforms $RPGR^{1\text{-}19}$ and $RPGR^{s14/15}$ in Wildtype, RPGR_KO2, and Δ14/15_2. *Vinculin* served as a loading control. Semi-quantitative densitometric measurements of the RT-PCR band intensities relative to *Vinculin* are shown in the right panel. **C.** Western blot analysis of RPGR isoforms $\mathrm{RPGR}^{1\text{-}19}$ and $\mathrm{RPGR}^{s14/15}$ in Wildtype, RPGR_KO2, and Δ14/15_2 cells. Orange asterisk indicates unspecific bands. GAPDH served as a loading control. Semi-quantitative densitometric analyses of the protein band intensities relative to GAPDH are presented in the right panel. One-way ANOVA using GraphPad Prism was used to calculate the *p* values (non-significant: $p \geq 0.05$; significant: $p < 0.01$).

### Table S1: Verification of possible off-target effects of the CRISPR approach.

One, two or three mismatches of the sgRNA with the genomic binding site (human genome assemble hg38) were considered. We analyzed possible sequence alterations at the potential sgRNA binding sites.

| **sgRNA *RPGR*-int13: atgtatcacagactagagagTGG** | | |
|---|---|---|
| **genomic loci** | **number of mismatches** | **sequence status** |
| Chr3:104917303 | 3 | reference sequence |
| Chr4:65425111 | 3 | reference sequence |
| Chr4:94500518 | 3 | reference sequence |
| Chr10:107931947 | 3 | reference sequence |
| Chr11:66644278 | 3 | reference sequence |
| Chr14:72332247 | 3 | reference sequence |
| **sgRNA *RPGR*-int15: cagtacatttggttagttagGGG** | | |
| **genomic loci** | **number of mismatches** | **sequence status** |
| Chr2:130374029 | 3 | reference sequence |
| Chr4:48616944 | 3 | reference sequence |
| Chr6:80072108 | 3 | reference sequence |
| Chr8:126451638 | 3 | reference sequence |
| Chr15:69180148 | 3 | reference sequence |
| Chr18:63987989 | 3 | reference sequence |
| ChrX:143010616 | 3 | reference sequence |
| ChrX:12647599 | 3 | reference sequence |
| **sgRNA *RPGR*-exon2: ctccacatgaaagatgtacaGGG** | | |
| **genomic loci** | **number of mismatches** | **sequence status** |
| chr2:133668243 | 3 | reference sequence |
| chr2:214553383 | 3 | reference sequence |
| chr5:179442148 | 3 | reference sequence |
| chr7:138500083 | 3 | reference sequence |
| chr8:141684357 | 3 | reference sequence |
| chr8:144377698 | 3 | reference sequence |
| chr15:91262748 | 3 | reference sequence |
| chr17:4452383 | 2 | reference sequence |
| chr17:69363488 | 3 | reference sequence |
| chr19:13171258 | 3 | reference sequence |
| chr21:41529628 | 3 | reference sequence |
| chrX:712431 | 3 | reference sequence |
| chrY:712431 | 3 | reference sequence |

**Table S2: Primer sequences and PCR conditions.**

| Application | Primer sequence | PCR conditions | |
|---|---|---|---|
| Genotyping of RPGR_KO1 and RPGR_KO2 | Forward: 5′- AGAAGGAAGGCTTAAACATTGC-3′<br>Reverse: 5′- TTCATTCCAAGAAAGTTGTGTGT-3′ | 95º C 15 min<br>95º C 45 sec<br>63º C 45 sec<br>72º C 35 sec<br>72º C 10 min | 32x |
| Genotyping of Δ14/15_1 and Δ14/15_2 | Forward: 5′- TGGCAGGTAGTAAGAATCGAAA-3′<br>Reverse: 5′- CTAGGGAGGCCAGTGTTCTC-3′ | | |
| *RPGR* transcript analysis between exon 1 and exon 5 | Forward: 5′- GCATGAGGGAGCCGGAAGAGC-3′<br>Reverse: 5′- TGTCCTTCATTATTTCCACCAG-3′ | 95º C 15 min<br>95º C 45 sec<br>62º C 45 sec<br>72º C 35 sec<br>72º C 10 min | 32x |
| *RPGR* transcript analysis between exon 11 and exon 16 | Forward: 5′- GGGACTCTTGGCCTTTCTGCTTGTT-3′<br>Reverse: 5′- TTTCAGCATTAATTTCCTCATCCACATCT-3′ | | |
| Vinculin transcript analysis | Forward: 5′- AGAGAAGCCTTCCAACCTCAG-3′<br>Reverse: 5′- CCTTCTGCTCAGGGAACTCTT-3′ | 95º C 15 min<br>95º C 45 sec<br>62º C 45 sec<br>72º C 35 sec<br>72º C 8 min | 28x |

**Table S3: Antibodies for immunocytochemistry (ICC) and Western blot (WB) analysis.**

| Antibody | Manufacturer | Catalogue number | Use | Dilution (ICC/WB) |
|---|---|---|---|---|
| Arl13B | Proteintech | 17711-1-AP | ICC | ICC-1:500 |
| GT335 | Biomol | AG-20B-0020-C100 | ICC | ICC-1:500 |
| RPGR | Sigma | HPA001593 | ICC/WB | ICC-1:250/ WB-1:500 |
| GAPDH | Chemicon Int. (Merck) | mab374 | WB | WB-1:1000 |
| Alexa Fluor™ 647 Phalloidin | Thermo Fischer Scientific | A22287 | ICC | ICC-1:250 |
| IgG anti-mouse Alexa Fluor™ 488 | Life Technologies | A21202 | ICC | ICC-1:1500 |
| IgG anti-rabbit Alexa Fluor™ 568 | Life Technologies | A10037 | ICC | ICC-1:1500 |
